# Familiarity modulates preference for joint feeding with conspecifics in rats

**DOI:** 10.1101/2024.04.18.590018

**Authors:** Reo Wada, Makiko Kamijo, Noriko Katsu, Shiomi Hakataya, Kazuo Okanoya, Hiroki Koda

## Abstract

Rats are highly social mammals that form social groups of various sizes. Interestingly, they are known to show prosocial behaviors, and are socially tolerant of other conspecifics as they feed together, but the specificity of their prosociality has not been fully understood. Here, we investigated what degree rats showed social choice, i.e., preferential joint feeding even with unfamiliar animals by performing a simple feeding site choice task. In the task, subjects were tested to choose between the feeding site where they fed with other rats (social option) or the other site where they could feed alone (solitary option). The results showed that social option choices increased particularly when the partner was a single non-cagemate rat. This result means that high social preference for other individuals occurs spontaneously, even when they are feeding with unfamiliar non-cagemates. This high degree of social tolerance would lead to the suppression of aggressive behavior as well as enhancement of affiliative relationships even in situations when the rats met with unfamiliar conspecifics for the first time.

**Highlights:**

- Rats are highly socially tolerant mammals as they feed with conspecifics together.
- We explored how the presence of conspecifics affects feeding site choices in rats.
- Social choices increased when the partners were non-cagemates.
- Social choices decreased as the number of non-cagemates increased.
- Joint feeding with a single non-cagemate is observed particularly frequently.

## 1. Introduction

Rats are laboratory mammals derived from wild Norway rats, *Rattus norvegicus*. They developed high sociality through domestication (Schweinfurth, 2020; Wu & Hong, 2022). Interestingly, competitive or aggressive inter-individual conflicts among conspecifics are rarely observed in rats even in foraging (Weiss et al., 2017), but rather, they followed and feed with conspecifics together (Alfaro & Cabrera, 2021; Takahashi et al., 2015; Weiss et al., 2017). This is consistent with the reported high prosociality of rats (Bartal et al., 2011), indicated by rats’ high social preference and spontaneous helping behavior toward other individuals.

Joint feeding, especially by multiple animals in close proximity, is not so commonly observed in all animals living in groups (e.g., Jaeggi & Gurven, 2013; Rapaport & Brown, 2008) due to an ecological disadvantage of resource competition among individuals. Similarly, food sharing has been studied as a rare social behavior (Stevens & Gilby, 2004). To share the same food, the aggressive impulsivity resulting from the competitive relationship must be relaxed: animals must be tolerant of others. Prosocial animals generally show social tolerance, suggesting a link between the two (Cronin, 2012). Joint feeding, which can be easily observed in rats (Weiss et al., 2017), may be explained by their reduced aggressive impulsivity toward other individuals, or may be the result of a remarkably increased preference for other individuals.

The purpose of this study was to examine, by means of behavioral experiments, how the rat’s prosocial traits influence its choice of joint feeding and how its choice decision differs depending on the presence of multiple familiar or unfamiliar conspecifics. Rats are known to have preference for unfamiliar other individuals (Hackenberg et al., 2021; Smith et al., 2015), and their various social behaviors can be altered by familiarity with other individuals (Alberts & Galef, 1973; Bartal et al., 2014; Kiyokawa et al., 2014; Rogers-Carter et al., 2018). However, previous studies have not considered familiarity with other individuals as a factor modulating joint feeding, which is one of the social behaviors (Alfaro & Cabrera, 2021; Tan & Hackenberg, 2012; Weiss et al., 2017). In addition, while feeding with multiple individuals can be a competitive situation, the greater the number of other individuals in a foraging area, the greater the proportion of rats’ behavior that follows and feeds with other individuals rather than feeding alone (Alfaro & Cabrera, 2021). Therefore, familiarity with other individuals and the number of individuals feeding together may influence the preference for joint feeding.

Here we performed a feeding site choice test in which rats were required to choose one of the two feeding sites (joint feeding or solitary feeding) and examined under what conditions they would choose joint feeding with other individuals. In this feeding site choice test, the animals were rewarded equally at both feeding sites, but were given the choice of “sharing” one feeding site with other individuals and feeding simultaneously (social choice/joint feeding choice) or feeding alone at the other feeding site (solitary feeding choice). Particularly, we tested if the preference would alter depending on whether the shared individuals were familiar or unfamiliar. Additionally, the effect of the number of other individuals sharing the joint feeding area was also examined.

## 2. Materials and methods

For details, see supplementary information. Experiments were reviewed and approved in advance by the Animal Experiment Ethical Committee, the University of Tokyo, Graduate School of Arts and Sciences, (#2022-5).

### 2.1 Animals and apparatus

Six 8-10 month old female Sprague-Dawley rats, kept as a cagemate group, were used as subjects. In addition, another six female rats kept as a group were used as unfamiliar non-cagemate individuals. We used a custom-made maze (Figure 1), consisted of three main areas: the center (the corridor, which served as the “start location”), left (“social option location”) and right (“solitary option location”) compartments, separated by the doors which slid up and down.

**Figure 1.**
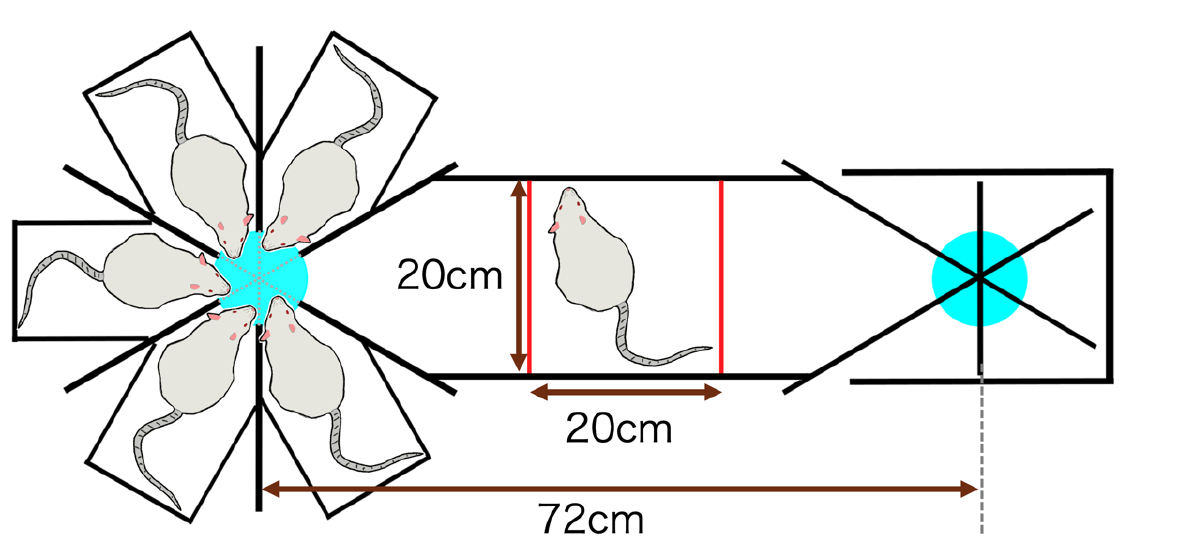
Schematic illustration of the apparatus. In the middle of the figure is the corridor, which served as the starting position, and the target subject animal was initially placed in this location. On the left is the social option, where the rats were placed in a small radial space. On the right side is the solitary option, a space of the same shape but without rats placed in it. The red lines represent doors, which slide up and down. Light blue circles are feeding dishes. In the experiment, experimental individuals obtain the same amount of food reward in both the social and the solitary options.

### 2.2 Procedure summary

The experiment consisted of four phases: a device habituation, a pretraining, an unbiased food search training, and a feeding site choice test (main test). The three phases conducted before the main test were performed to habituate the subjects to the apparatus and maze selection behavior and to remove positional bias. Following those three phases, rats were tested to choose one of two feeding sites (joint feeding/social option or solitary feeding/solitary option) in the maze (feeding site choice test). Each rat acquired an equal amount of food rewards (rice puffs) at either feeding site. Basically, twelve trials to choose sites were conducted, making one session. The experiment was conducted for two sessions per rat. The following factors and conditions were set for the introduced individuals: 1) cohabitation (cagemate or non-cagemate), and 2) number of other individuals (1, 3, or 5 individuals). These 6 conditions (cagemate/non-cagemate × 1/3/5 rats) were set, with the order of the conditions set randomly.

### 2.3 Analysis

We statistically tested by fitting with the generalized linear mixed model (GLMM), with a stepwise model selection procedure implemented by R version 4.3.2 (R Core Team, 2023).

## 3. Results

The model selection procedure reported the “full model”, including all explanatory terms, as the minimal AIC models (Table S1). The parameter coefficient of “cohabitation” was estimated as negative value (*β* ± *SE* = -1.431±0.442, *z* = -3.234, *p* = 0.001, the intercept of the model was set to “cagemate cohabiting condition”, see Table S2), that of “number of other individuals” was estimated to be not different from 0 (*β* ± *SE* = -0.107±0.086, *z* = -1.195, *p* = 0.232). The interaction effect term was estimated as positive value with statistical significance (β± *SE* = 0.320±0.127, *z* = 2.526, *p* = 0.012; Table S2). The results indicated that 1) social choices increased when the partners in the experiment were non-cagemate individuals, but 2) social choices decreased as the number of non-cagemate individuals increased (Figure 2). In other words, it suggested that rats chose the social option, particularly when one non-cagemate rat was placed on the social option side.

**Figure 2.**
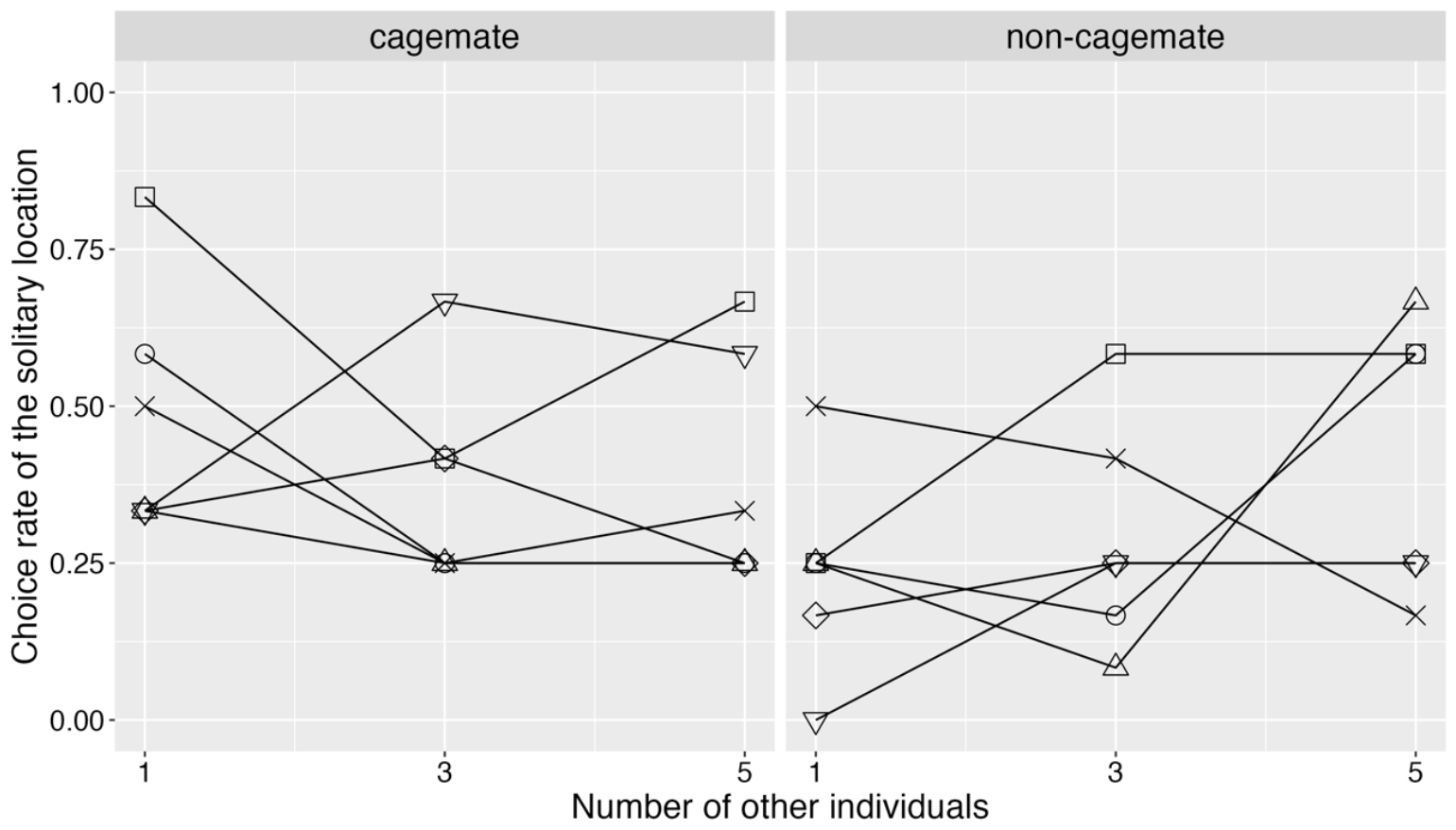
Plots of the choice of joint feeding/social and solitary options of the 6 rats for each condition. Each symbol represents the average of each individual’s choice score.

## 4. Discussions

In summary, the number of social option choices was more likely to increase when only one non-cagemate individual was placed on the social option side. At the same time, this effect was not observed in the case of cagemate individuals. These results suggest that this was due to a social preference, particularly for unfamiliar non-cagemate individuals with whom they do not usually have contact.

The social preference for unfamiliar individuals is consistent with the previous studies on strong preferences for non-cagemate or novel individuals (Alfaro & Cabrera, 2021; Hackenberg et al., 2021; Smith et al., 2015; Weiss et al., 2017). Rats often showed a novelty exploratory behavior (Berlyne, 1950), in which they search for novel objects rather than for familiar objects, suggesting that novelty increases social preference. Our results - higher preference for unfamiliar individuals than for familiar individuals – are due to the integration of novelty preference and social preference in rats, which may characterize the rats’ uniqueness of high prosociality.

It is still unclear why this social novelty preference is highest when there is only one other individual and weakens as the number of individuals increases. A simple interpretation is that approach to the social option side with multiple non-cagemates may be similar to the situation in which they approach the territory of another group. In fact, exclusive territorial defense behavior is known in rat groups (Blanchard et al., 1988). In contrast, a simple social novelty preference may have been observed in the one-individual situation, as the formation of territories does not firmly occur. In the case of cagemates placed on the social option, the avoidance of the social option may not have been enhanced by the increase in the number of cagemates, since they share the same territory.

Our experimental results reflect the social affiliation and prosociality of rats. Humans are unique animals that have acquired prosociality or social tolerance. These traits would be interpreted as adaptations to avoid aggression and to maintain higher cooperative groups (Burkart et al., 2014). Such an evolutionary process, involving selection for increased social tolerance toward other individuals, may bear some similarities in rats. Laboratory rats are highly domesticated, which may have increased their social tolerance, as they were initially adapted as pet animals. There is much evidence suggesting a coupling observation between domestication and high social tolerance, in humans, dogs, and other animals (Wilkins et al., 2014). The high social tolerance observed in rats, phylogenetically distant from humans or dogs, may suggest the possibility of convergent evolution as an adaptive characteristic to group living.

## Acknowledgements

We appreciate Noriko Kondo, Sota Kikuchi, Genta Toya. The study was supported by the Japan Society for the Promotion of Science Grant‐in‐Aid for Scientific Research (S, #23H05428) to KO, (A, #21H04421) to HK, Young Scientist (#21K13746) to MK, and JSPS Fellows (#22KJ0702) to SH, and by World-leading Innovative Graduate Study Program of Advanced Basic Science Course (WINGS-ABC) to RW.

