## supplementary information for "Familiarity modulates preference for joint feeding with conspecifics in rats"

### Supplementary methods

#### Animal details

The 12 female Sprague-Dawley rats were kept in separate cages of six rats each. Rats in one cage were used as test subjects, while rats in the other cage were used as “unfamiliar non-cagemates”. The housing environment was controlled with 12L:12D daylight, room temperature between 20-26°C, and humidity between 40-60%. They were fed daily with a fixed amount (app. 15g/individual) of pellets and allowed to drink water freely. The animals were habituated to the experimenter by handling them for 10 minutes per day per an animal during the week prior to the start of the experiment.

#### Apparatus details

We used a custom-made maze in which the subject rats had to choose between two locations to acquire food (Figure 1). The maze consisted of three main areas: the central area was the arm corridor, which served as the “start location”, the left side was the “social option location”, the right side was the “solitary option location”. The doors were placed between the start location and the choice locations on each side, which slid up and down. Detailed maze sizes and shapes were shown in Figure 1.

#### Procedures details

The experiment consisted of four phases: a device habituation, a pretraining, an unbiased food search training, and a feeding site choice test.

*Device habituation, pretraining, and unbiased food search training*

In the device habituation phase, all rats were allowed to explore freely in the apparatus to familiarize them with the corridor, the doors, and the door opening/closing operation. In the pretraining phase, they were placed in the start location for 10 seconds, and after the doors were opened, they were fed when they reached the food plate within 2 minutes to train their feeding behavior at the choice locations.

Next, the “unbiased food search training” was conducted to eliminate the left-right positional bias. This training consisted of two phases: a training phase and a positional bias confirmation test. In the training phase, after the rats waited in the start location for 10 seconds, the door was opened in a random order on either the left or right side, and the rats were fed with two rice puffs when they reached the end of the corridor. This was repeated for 24 trials (left: 12 trials, right: 12 trials). Following the training phase, the positional bias confirmation test was conducted. In this phase, the left and right doors were opened at the same time and the animals were allowed to freely select the left or right feeding site to examine the positional bias of the left or right. The positional bias confirmation test was conducted for 12 consecutive trials, and if 1) either side was selected between five and seven times, and 2) the last four trials were not in the same location side, the individual was considered to have no positional bias, and was transferred to the next phase of the experiment (Feeding site choice test). If positional bias was identified, the “unbiased food search training” was repeated until it was judged that there was no positional bias. Finally, it was confirmed that all rats had no positional bias.

#### *Feeding site choice test*

After the subject was confirmed to have no positional bias, the main test of the experiment, “feeding site choice test”, was conducted. Before starting the test, we introduced other individuals to the social option location. At the beginning of the test, we always started with 4 trials of the unbiased food search training. That is, the four trials (joint choice: 2 trials, solitary choice: 2 trials) were forced to be experienced in a random order. The two doors were then opened simultaneously and a free-choice trials were conducted.

In the free-choice trials, to avoid left-right positional bias being acquired during the test, a positional bias cancellation trial was inserted to force the subject to choose the opposite option if she chose the same option for two consecutive trials (e.g., if the subject rat chose the social option for two consecutive trials, she was forced to choose the solitary option once). Twelve free-choice trials were conducted with the insertion of the positional bias cancellation trial, making one session. The experiment was conducted for two sessions per rat.

The following factors and conditions were set for the introduced individuals: 1) cohabitation (cagemate or non-cagemate), and 2) number of other individuals (1, 3, or 5 individuals). These 6 conditions (cagemate/non-cagemate × 1/3/5 rats) were set, with the order of the conditions set randomly. After the test subject completed the feeding site choice test for one experimental condition, she underwent the unbiased food search training again before moving on to another condition.

#### Analysis

To statistically test whether cohabiting conditions and number of individuals affected the choice of options, we statistically assessed the results by fitting with the generalized linear mixed model (GLMM), where the model formula was set to "glmer(choice ~ cohabitation * number + (1|subject), family = binomial)", accounting for the binary choice data (0 := joint feeding/social option and 1 := solitary feeding/solitary option), in "glmer()" function of the "lme4" package in R. Next, the "best" model, i.e., the minimum AIC model, was estimated by a stepwise model selection procedure implemented by by "deglege()" in the R package "MuMIn". Finally, we report the best model and the estimated parameter coefficients, and those 95 % confidence intervals by "confint" function in "stats" package.

### Author contributions

**Reo Wada**: Conceptualization, Methodology, Formal analysis, Investigation, Writing - Original Draft, Visualization. **Makiko Kamijo**: Conceptualization, Writing - Review & Editing, Funding acquisition. **Noriko Katsu**: Conceptualization, Writing - Review & Editing. **Shiomi Hakataya**: Conceptualization, Methodology, Writing - Review & Editing. **Kazuo Okanoya**: Conceptualization, Resources, Writing - Review & Editing, Supervision, Funding acquisition. **Hiroki Koda**: Conceptualization, Resources, Investigation, Writing - Original Draft, Supervision, Funding acquisition.
