## Supplementary material for "Familiarity modulates preference for joint feeding with conspecifics in rats": Table S1

Table S1. Model selection table sorted by AIC.

| Model order | Cohabiting(cagemate or non-cagemate) | Number of individuals | Cohabiting:Number of individuals | Df | AIC | Δ AIC |
|---|---|---|---|---|---|---|
| 1 | * | 0.217 | * | 5 | 562.0 | |
| 2 | * | | | 3 | 565.0 | 3.03 |
| 3 | * | 0.046 | | 4 | 566.4 | 4.48 |
| 4 | | | | 2 | 567.9 | 5.98 |
| 5 | | 0.046 | | 3 | 569.4 | 7.43 |

Results of the model selection procedure for the GLMM accounting for the solitary option selection. The models are listed in ascending order, from the model with the smallest AIC (the best model) to the model with the largest AIC. Asterisks (*) indicate explanatory variables included in the model, and the differences between the AIC of the model and the model with the smallest AIC are shown as ΔAIC.
