## Supplementary material for "Familiarity modulates preference for joint feeding with conspecifics in rats": Table S2

Table S2. Parameter coefficients of the best GLMM for the binary data of social/solitary options.

| | estimate | SE | z value | Pr (>\|z\|) | 95% CI |
|---|---|---|---|---|---|
| Intercept | -0.037 | 0.316 | -0.118 | 0.906 | [-0.668, 0.591] |
| Cohabiting (A) | -1.431 | 0.442 | -3.234 | 0.001 | [-2.314, -0.574] |
| # individuals (B) | -0.107 | 0.086 | -1.195 | 0.232 | [-0.272, 0.065] |
| A x B | 0.320 | 0.127 | 2.526 | 0.012 | [0.073, 0.571] |

Intercept of the model was set to the “cagemate cohabiting condition.” Results of the parameter coefficients of the best GLMM. The 95 % confidence intervals were computed in the best model by “confint” function of R.
